# Deep Learning Enhances Precision of Citrullination Identification in Human and Plant Tissue Proteomes

**DOI:** 10.1101/2024.10.11.617769

**Authors:** Wassim Gabriel, Rebecca Meelker Gonzalez, Sophia Laposchan, Erik Riedel, Gönül Dündar, Brigitte Poppenberger, Mathias Wilhelm, Chien-Yun Lee

## Abstract

Citrullination is a critical yet understudied post-translational modification (PTM) implicated in various biological processes. Exploring its role in health and disease requires a comprehensive understanding of the prevalence of this PTM at a proteome-wide scale. Although mass spectrometry has enabled the identification of citrullination sites in complex biological samples, it faces significant challenges, including limited enrichment tools and a high rate of false positives due to the identical mass with deamidation (+0.9840 Da) and errors in monoisotopic ion selection. These issues often necessitate manual spectrum inspection, reducing throughput in large-scale studies. In this work, we present a novel data analysis pipeline that incorporates the deep learning model Prosit-Cit into the MS database search workflow to improve both the sensitivity and precision of citrullination site identification. Prosit-Cit, an extension of the existing Prosit model, has been trained on ∼53,000 spectra from ∼2,500 synthetic citrullinated peptides and provides precise predictions for chromatographic retention time and fragment ion intensities of both citrullinated and deamidated peptides. This enhances the accuracy of identification and reduces false positives. Our pipeline demonstrated high precision on the evaluation dataset, recovering the majority of known citrullination sites in human tissue proteomes and improving sensitivity by identifying up to 14 times more citrullinated sites. Sequence motif analysis revealed consistency with previously reported findings, validating the reliability of our approach. Furthermore, extending the pipeline to a tissue proteome dataset of the model plant *Arabidopsis thaliana* enabled the identification of ∼200 citrullination sites across 169 proteins from 30 tissues, representing the first large-scale citrullination mapping in plants. This pipeline can be seamlessly applied to existing proteomics datasets, offering a robust tool for advancing biological discoveries and deepening our understanding of protein citrullination across species.

## Introduction

Citrullination is a post-translational modification (PTM) of arginine residues catalyzed by peptidylarginine deiminases (PADs). This modification results in the loss of a positive charge and introduces a mass shift of +0.9840 Da, which can significantly impact the structure and function of the modified protein, potentially increasing immunogenicity or impairing binding interactions (1, 2). Citrullination has been linked to several diseases in humans, including rheumatoid arthritis (RA), cancers, and multiple sclerosis (2, 3). In other higher organisms, such as plants, the functional significance of this type of PTM has remained largely understudied (4).

Mass spectrometry (MS) has become a powerful tool for the large-scale study of protein PTMs, providing site-specific and quantitative information (5). In typical PTM proteomics workflows, proteins are digested into peptides, often using trypsin, followed by enrichment techniques to detect low-abundance PTMs (6). While several probes and antibodies are available for detecting citrullination, there are no pan-citrullination antibodies suitable for large-scale MS studies, and chemical probes often suffer from poor fragmentation efficiency during MS analysis (7, 8).

Without specific enrichment, low-abundant citrullination can still be identified through deep proteome profiling (9, 10). Techniques such as orthogonal separation using basic reverse-phase high-performance liquid chromatography (RP-HPLC) prior to MS analysis enable the identification of low-abundance modified peptides within datasets covering over 8,000–10,000 proteins (5, 11). However, two major challenges complicate citrullination identification: the identical mass shift between citrullination (Arg) and deamidation (Asn/Gln), leading to misidentification, and incorrect selection of monoisotopic precursors due to natural isotopic variants. Methods such as two-step database searching (12) and diagnostic ion mining (13) have been proposed to address these issues, but their sensitivity has not been fully demonstrated on large-scale proteomics datasets.

Moreover, controlling the false discovery rate (FDR) in large-scale proteomics datasets for rare PTMs like citrullination presents additional difficulties. The lower number of citrullination identifications often skews score distributions, complicating FDR optimization (14). Approaches such as “subset/group FDR” (14) or “subset-neighbor search” (SNS) (15), which apply FDR control to a limited set of identifications, or only to a defined set of identifications, have been used for PTM proteomes (16). However, none of these methods have been extensively tested for citrullination identifications.

Recent advances in deep learning have significantly improved the analysis of MS-based proteomics. These models enhance the discrimination between correct and incorrect peptide-spectrum matches (PSMs) by incorporating features such as retention time predictions and correlations between observed and predicted MS/MS (MS2) spectra (17–22). Among them, Prosit has been particularly successful in boosting peptide identification for various proteomics applications, including global profiling, immunopeptidomics, and isobaric labeling (17, 19, 21, 23, 24). However, none of the existing models, including Prosit, has been trained to predict fragment ion intensities for citrullinated peptides. Since citrullination alters fragmentation patterns (25), untrained models often result in inaccurate fragment ion predictions. To accurately predict the fragmentation patterns of citrullinated peptides, training on a substantial set of high-quality, synthetic peptide spectra is essential.

Here, we present a new data analysis pipeline (Figure 1A) that accurately identifies citrullination sites in proteomics datasets by incorporating Prosit-Cit, an extension of the Prosit framework (17, 19) tailored to citrullinated peptides. Prosit-Cit is a fine-tuned model trained additionally on ∼53,000 spectra from approximately 2,500 synthetic citrullinated peptides generated by the ProteomeTools project (26). This pipeline delivers high precision in citrullination identification, validated using 200 synthetic citrullinated peptides (25) spiked into human proteome digests. Applying this approach to the analysis of ten human tissue proteomes (9, 27) uncovered up to 14 times more citrullinated sites compared to the previous study. Extending this search to tissue proteomes of *Arabidopsis thaliana* (Arabidopsis) (28) revealed ∼200 novel citrullination sites from 169 proteins. This work provides a precise and high-throughput solution for large-scale citrullination mining and represents the first comprehensive survey of protein citrullination in plants, enabling deeper insights into the biological significance of this modification.

**Figure 1.**
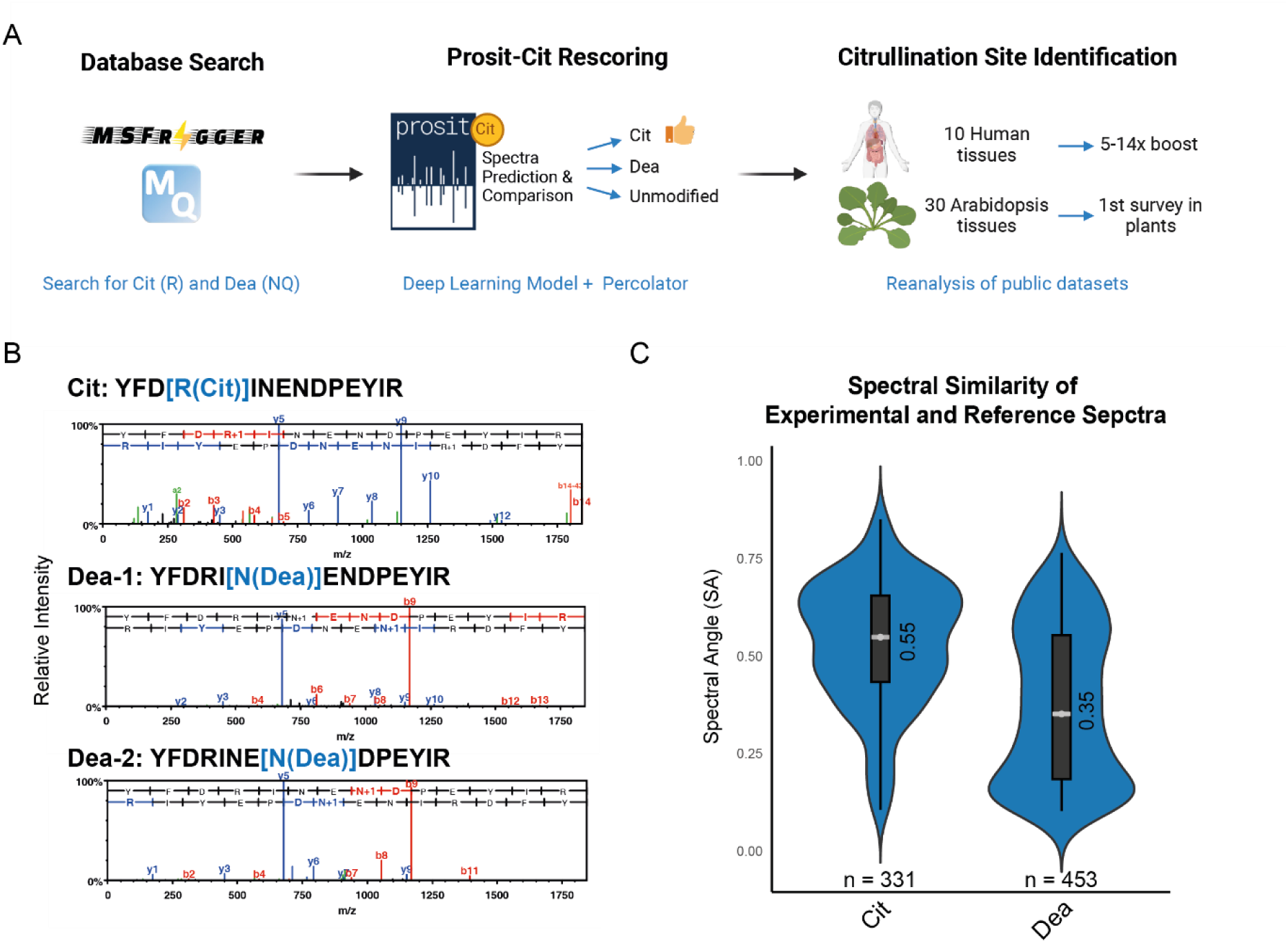
Deep learning boosts citrullination identification in human and Arabidopsis tissue proteomes. **A**. Workflow illustration. **B.** MS2 spectra of synthetic peptides containing citrullination or deamidation of gamma-adducin. **C.** Spectral angle comparison of experimental spectra with reference spectra of citrullination and deamidation from Lee *et al.* (9). Cit, citrullinated/citrullination; Dea, deamidated/deamidation.

## Experimental Procedures

### Experimental Design and Statistical Rationale

This study presents a data analysis workflow designed to improve the precision of citrullinated peptide identification using the deep learning framework Prosit-Cit. Detailed workflow and model generation descriptions are provided in subsequent sections, along with the results. Briefly, an evaluation dataset was constructed to assess the precision of various database search, rescoring, and post-processing steps. This dataset includes a complex biological matrix (MES-SA cell tryptic digest) spiked with approximately 200 synthetic citrullinated peptides (25) at varying ratios. After database searching and rescoring, the dataset was annotated with true and false positive identifications. This optimized workflow was then applied to two published proteomic datasets—human and Arabidopsis tissue proteomes—to explore citrullination sites.

### Synthetic Citrullinated Peptides

The synthetic peptides used in this study were part of the ProteomeTools project, and their selection and synthesis have been described in earlier studies (25, 26). In summary, all peptides were tryptic and synthesized via Fmoc-based solid-phase SPOT synthesis (29) (JPT Peptide Technologies GmbH, Germany). Two pools of synthetic citrullinated peptides were utilized. The first pool, used for spike-in experiments, contained 200 synthetic citrullinated peptides, each with one citrullination site (25). The second pool consisted of ∼2,500 citrullinated peptides previously reported in the literature, which may have more than one citrullination site per peptide (9).

### Cell Culture, Lysis, and Protein Digestion

The sarcoma cell line MES-SA was cultured in McCoy’s 5A Medium Modified, supplemented with 10% fetal bovine serum (FBS) (PAN Biotech GmbH, Germany). Cells were lysed in a buffer containing 40 mM Tris-HCl (pH 7.6) and 2% SDS, followed by boiling at 95°C for 5 minutes. Protein concentration was determined using the Pierce BCA Protein Assay Kit (Thermo Scientific, MA, USA). Lysates were processed using the SP3 method (30). In brief, 10 µL of a carboxylate bead mix (50 µg/µL in H₂O, Sera-Mag Speed beads, GE Healthcare, IL, USA) was added, followed by one precipitation step with 70% acetonitrile (ACN) and two wash steps with 80% ethanol, and one wash step with 100% ACN. The beads were then incubated in 90 µL digestion buffer (100 mM HEPES, pH 8.5, 10 mM DTT) at 37°C and 600 rpm for 1 hour. Subsequently, 10 µL of 550 mM chloroacetamide (CAA) was added, followed by a 30-minute incubation in the dark. After reduction and alkylation, the samples were digested overnight at 37°C and 800 rpm using a 1:50 trypsin-to-protein ratio. Digested peptides were desalted using a 50 mg Sep-Pak cartridge and dried for further analysis.

### Generation of Evaluation Dataset

The synthetic citrullinated peptide pool (25) was dissolved in 0.1% formic acid (FA) to a final stock concentration of 200 pmol/µL. Six dilutions were prepared from this stock using a 1:3 dilution series (200 to 0.27 pmol/µL). Four microliters of each stock and its dilution series were mixed with two µg of tryptic-digested MES-SA cell lysate for downstream analysis.

### Data Acquisition and Database Searching

The evaluation dataset was analyzed using nano-liquid chromatography coupled with an Orbitrap Lumos mass spectrometer (Thermo Fisher Scientific, Bremen, Germany) in a 60-minute gradient data-dependent acquisition (DDA) method (4-32% ACN). The MS settings included an Orbitrap full MS scan at 60,000 resolution (3e6 AGC target, 45 ms maximum injection time, 360–1,300 m/z, profile mode), followed by up to 18 MS/MS scans at 15,000 resolution (1e5 AGC target, 25 ms maximum injection time, 1.3 m/z isolation width, centroid mode, 26 normalized collision energy (NCE)) with a dynamic exclusion of 25 seconds. Five hundred nanograms of peptides were injected for each run.

Raw files from the evaluation dataset were searched using both MaxQuant (version 2.2.0.0) (31) and Fragpipe (version 20.0 with MSFragger v3.8, IonQuant v1.9.8, and Philosopher v5.0.0) (20, 32). Both searches were conducted against the UniProtKB-SwissProt complete human proteome (canonical version, downloaded on 03.01.2023) using default settings, with the following specific modifications: cysteine carbamidomethylation was set as a fixed modification, and variable modifications included methionine oxidation, arginine citrullination, asparagine and glutamine deamidation, and N-terminal acetylation. Strict trypsin digestion was applied, allowing for up to three missed cleavages. Precursor and fragment mass tolerances were set to 20 ppm (with 4.5 ppm for MaxQuant main searches). The mass calibration option was enabled in MSFragger. Both searches were conducted with a 1% FDR and 100% FDR at the PSM level for Prosit-Cit rescoring. Fragpipe searches were performed with and without MSBooster rescoring, and PSMs were validated using Percolator with ProteinProphet enabled. The search parameter files (mqpar.xml and fragpipe.workflow) can be accessed in the PRIDE repository with the dataset identifier PXD056560 along with the results.

### High-Quality LC-MS/MS Dataset from ProteomeTools

The reference spectra used in this study are part of the ProteomeTools project, a synthetic peptide library from the human proteome to advance biomedical research (26). Over 30 million tryptic and non-tryptic reference spectra have been published and used for training Prosit (17, 19). Spectra for each peptide were generated under various mass spectrometry conditions (25, 26). Citrullinated peptides from this study were analyzed using three methods: (i) a DDA method consisting of an Orbitrap full MS scan followed by HCD-based MS/MS scans; (ii) a modified DDA method with consecutive HCD scans at 20, 25, and 30 NCE; and (iii) a combination of MS1 scans with MS2 scans using CID or HCD fragmentation with ion trap or Orbitrap readouts. Spectra from the first method have been published (9), and the rest can be accessed in the PRIDE repository with the dataset identifier PXD056559 along with the search results.

### Deep Learning Using Prosit Framework

The architectural framework utilized in the Prosit model is aligned with the one specified by Gabriel *et al.* (24). Regarding the intensity model, the training was performed as described by Wilhelm *et al.* (19)and as described by Gessulat *et al.* (17) for the retention time model. Raw files were converted to mzml format utilizing the Thermo rawfile parser (https://github.com/compomics/ThermoRawFileParser) to facilitate the extraction of MS2 spectra. The spectra were annotated employing b- and y-ion coverage for fragment charges ranging from 1 to 3, ensuring that the fragment charge did not exceed the observed precursor charge. Annotation of spectra obtained through FTMS and ITMS readout was executed with a tolerance threshold of 20 ppm and 0.4 Da, respectively. Shouman *et al.* (33) articulate additional details regarding the data processing. Peptides exhibiting lengths less than 7 or exceeding 30, with precursor charges greater than 6, and PSMs with an Andromeda score below 60 were excluded from the training dataset. The five highest-scoring PSMs for each respective peptide were utilized for the training procedure. The data was partitioned at the unmodified peptide level into 70% for training (∼12.1M spectra), 20% for validation (∼3.5M spectra), and 10% for testing (∼1.7M spectra). The data splitting was done randomly while guaranteeing that each peptide sequence was confined to one of the datasets. Annotated spectra accompanied by metadata are archived as parquet files in Zenodo (https://zenodo.org/records/13856705).

### Rescoring Pipeline

The following steps were used to rescore and localize citrullination and deamidation identifications. The pipeline tutorial (Prosit_cit_tutorial) can be accessed via GitHub (https://github.com/wilhelm-lab/oktoberfest/tree/development/tutorials).

#### Preprocessing

The search results with 100% FDR from MaxQuant or MSFragger were used. For all peptides with either citrullination (cit) or deamidation (dea) identified, the “neighbor” for this peptide sequence were produced, where the other potential modifications of +0.98 Da on R, Q, N of the sequence may occur. Besides, the unmodified counterpart was also generated. For instance, if a citrullination was identified in PEPNTIDQE[R(cit)]PEK, the combinations of the following will be generated: 1) PEPNTIDQE[R(cit)]PEK, 2) PEPNTID[Q(dea)]ERPEK, 3) PEP[N(dea)]TIDQERPEK, and 4) PEPNTIDQERPEK.

#### Additional scores

For all citrullinated and deamidated PSMs and their combinations, distinct scores were computed from predictions, as described by Gessault *et al.* (17). Two additional scores were incorporated to refine the scoring of the citrullinated and deamidated variants: (1) Neutral Loss Peaks: A score was calculated based on the number of peaks exhibiting isocyanic acid (CHNO) neutral loss from fragment ions containing citrullination. (2) Theoretical Neutral Loss: A count of theoretically feasible neutral loss peaks in the spectrum, based on the modified peptide sequence.

#### First Percolator run

We then ran Percolator (34) on this set of PSMs, where the citrullinated and deamidated variants comprised the relevant set, and their complementary and unmodified counterparts formed the neighbor set, following the subset-neighbor search (SNS) approach (15). This method ensures that both the modified and unmodified variants are compared together in order to distinguish modifications confidently. From the first Percolator run, the highest-scoring match for each scan was selected. Any scan where the unmodified peptide was chosen as the best match was excluded from further analysis.

#### Second Percolator run

The second Percolator run was applied only to the citrullinated and deamidated PSMs from the first run, following a method analogous to the group FDR approach (14). In this step, a 1% FDR threshold was applied at the PSM level.

### Reanalysis of External Datasets

Raw files of the human (28) and Arabidopsis (28) tissue proteomes were downloaded from PRIDE and searched using MSFragger against UniProtKB-Swiss-Prot human and Arabidopsis proteomes (downloaded from Uniprot on January 3, 2023, and January 5, 2024, respectively). Searches were conducted with the previously mentioned parameters, with an initial 100% FDR. These results were rescored using Prosit-Cit and Percolator, refining the FDR to 1% at the PSM level. Target files (.txt) from these searches were annotated with site and sequence information using a Python package (https://github.com/kusterlab/psite_annotation). Quantification of PSMs from the target text files was matched to the corresponding modified peptides from Fragpipe (IonQuant) using raw file names and scan numbers. The annotation step is included in “Prosit_cit_tutorial” available on GitHub (https://github.com/wilhelm-lab/oktoberfest/tree/development/tutorials).

### Motif Analysis

Potential substrate motifs surrounding identified citrullination sites were analyzed using pLOGO (35). Sequences spanning −5 to +5 residues around the citrullination sites were extracted and compared with their respective background frequencies in the human or Arabidopsis proteomes. Significant motifs (*p* = 0.05, Bonferroni corrected) were visualized.

### Growing Conditions, Cold Treatment, and Protein Extraction of Arabidopsis Flower Tissues

Arabidopsis plants of the ecotype Columbia-0 (Col-0) were grown in soil for 5 weeks in long-day growth conditions (16 hrs of white light at 80 μmol m−2 s−1/ 8 hrs of dark; in a BrightBoy growth chamber, CLF Plant Climatics) at 22 °C +/− 1° to the flowering stage. The plants were then subjected to a cold treatment of 4 °C for either 90 min or 24 h using a controlled temperature chamber (Panasonic MIR-154, Panasonic Biomedical), and flowers were harvested immediately after the treatment. For each treatment, five individual plants were used and the samples were pooled. Protein extraction was carried out after grinding the plant material, adding 500 μL of protein extraction buffer (50 mM NaCl, 50 mM Tris-HCl, pH 7.5, 1 mM EDTA, 1% Triton X-100, 1 mM DTT, 1 mM PMSF) and centrifugation at 13,300 rpm for 15 minutes at 4°C. The resulting supernatant was collected, and protein concentrations were determined using the Pierce BCA Protein Assay Kit (Thermo Scientific, MA, USA).

### Flower Sample Preparation and Mass Spectrometry Analysis

Protein extracts from Arabidopsis flower samples underwent a two-step protein precipitation protocol (36). The resulting protein pellet was processed using the SP3 method (30), followed by tryptic digestion overnight. About 75 micrograms of digested peptides were desalted and fractionated using self-constructed C18 StageTips (C18 extraction disc, IVA Analysentechnik GmbH, Germany) as previously described (37). After acidifying the peptide digests with FA, the peptides bound to the StageTips were washed with 0.1% FA and then equilibrated with 25 mM ammonium formate (pH 10). Sequential elution of peptides was performed using increasing concentrations of acetonitrile (5%, 7.5%, 10%, 12.5%, 15%, 17.5%, 50%) in 25 mM ammonium formate. Six fractions were collected, with the desalted flow-through fraction combined with the 17.5% ACN fraction and the 5% ACN fraction combined with the 50% ACN fraction. All fractions were acidified with formic acid to a final concentration of 1%, dried, and stored at −20°C until MS analysis. For LC-MS/MS, peptides were resuspended in 0.1% formic acid and analyzed on an Orbitrap HF-X mass spectrometer (Thermo Fisher Scientific, Bremen, Germany) using similar settings as the evaluation dataset. Each injection consisted of 500 ng of peptides. Raw files were searched against the UniProtKB-Swiss-Prot Arabidopsis proteome (downloaded on January 5, 2024), with rescoring by Prosit-Cit and Percolator at 1% FDR on the PSM level.

### Western Blotting

Protein samples (40 μg) from flowers were separated via SDS-PAGE and transferred onto PVDF membranes. Membranes were blocked with 2% BSA in TBS-T (0.05% Tween-20 in Tris-buffered saline) for 1 hour at room temperature. Following blocking, the membranes were incubated overnight at 4°C with mouse anti-citrullinated peptide antibody clone F95 (MABN328; MilliporeSigma, MA, USA). After washing with TBS-T, the membranes were incubated with a goat anti-mouse IgM secondary antibody conjugated to a fluorophore (LI-COR Biosciences GmbH, Germany). Signals were detected using the Odyssey imaging system (LI-COR Biosciences GmbH, Germany). For loading control, the SDS-PAGE were stained with the Coomassie Brilliant Blue dye.

## Results

### MS2 Spectra Discriminate Citrullination from Deamidation

Neutral loss (NL) ions from isocyanic acid serve as reliable diagnostic markers for citrullinated peptides, notably improving the accuracy of citrullination site identification (9, 38). However, the so-called “citrullination effect” (39), characterized by a cleavage bias at the C-terminus of citrulline residues, results in low-abundant citrulline-containing y ions (∼44%). This often leads to the absence of NL ions in spectra, complicating accurate peptide identification. Previous research demonstrated that citrullinated peptides exhibit unique MS2 spectra compared to their unmodified counterparts (25), enabling better discrimination. Yet, the similarity between the spectra of citrullinated and deamidated peptides remains largely uncharacterized and requires further exploration.

To evaluate whether b- and y-ion intensities in MS2 spectra can reliably differentiate citrullinated peptides from deamidated ones, we analyzed a synthetic peptide spectral library (HCD) generated in the previous study (9). This library consists of over 2,200 citrullinated peptides and 1,300 deamidated peptides, where deamidated Asn and Gln residues were replaced by Asp and Glu, respectively. A notable distinction in fragment ion intensities was observed between citrullinated and deamidated peptides. For instance, a peptide from Gamma-adducin (Figure 1B) displayed a difference in spectral intensities between its citrullinated and deamidated forms. While both peptides shared an abundant y5 ion, the citrullinated peptide exhibited higher intensities in the y6–y10 ion series, which were absent in the deamidated spectra. Additionally, the “citrullination effect,” characterized by diminished y11–y13 ions, rendered the NL undetectable in the citrullinated peptide—a phenomenon not seen in deamidated peptides. We further analyzed spectral angle (SA) scores to assess the similarity between MS2 HCD spectra from experimental and synthetic peptides, both citrullinated and deamidated, based on comparisons from the same study (9). SA scores are indicative of spectral similarity, with higher scores reflecting greater alignment (40). Spectral comparisons from 784 synthetic peptides and 394 experimental spectra (manually validated as citrullinated) (Figure 1C; Supplementary Table S1) demonstrated high SA scores when citrullinated reference spectra were compared, but lower SA values with deamidated reference spectra. These observations suggest the ability of MS2 spectra to discriminate citrullinated from deamidated peptides.

### Evaluating Search Engine Performance in Citrullination Identification Precision Using a Custom Dataset

As the potential of MS2 spectra to discriminate between citrullinated, deamidated, and unmodified peptides was demonstrated, we hypothesized that incorporating spectral comparison methods into database search pipelines would improve the precision of citrullination identification. To evaluate this, we first generated an evaluation dataset by spiking seven different concentrations of 200 synthetic citrullinated peptides from the ProteomeTools project (25) into tryptic digests of the human proteome. The peptide concentrations ranged from 1 fmol to 1,000 fmol, simulating biological sample conditions (Figure 2A). These samples were analyzed using a 1-hour gradient on LC-Orbitrap MS/MS with HCD fragmentation. The resulting dataset was used to assess the precision of citrullination identification across various proteomics search engines.

**Figure 2.**
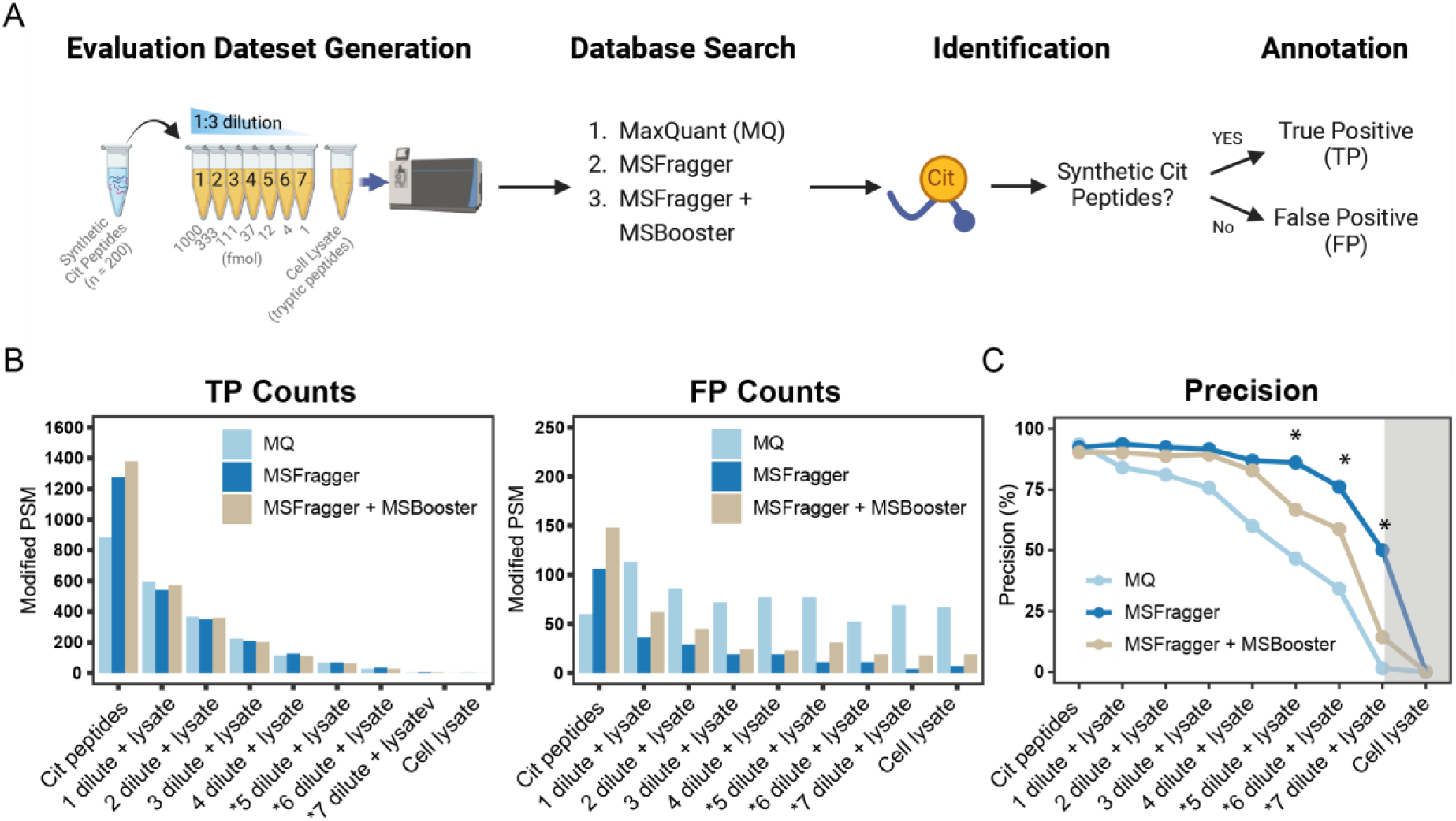
Comparing the precision of citrullination identification in MaxQuant and MSFragger using the evaluation dataset. **A.** Annotation of true positives and false positives in the evaluation dataset. **B.** True and false positive counts of modified peptide-spectrum matches (PSMs). **C.** Precision of citrullination identification, calculated as TP/(TP+FP). MQ, MaxQuant. Cit, citrullinated. TP, True positive; FP, False positive. Asterisk indicates the range of citrullination level in human tissue proteomes (9).

The dataset was searched using MaxQuant (MQ) (31) and MSFragger (32), both with citrullination and deamidation specified as variable modifications. These engines were selected due to their academic accessibility and robust quantification features suitable for large-scale proteomics. Additionally, MSBooster was enabled in MSFragger to enhance peptide identification by rescoring peptide spectrum matches (PSMs) using deep learning-based predictions of peptide properties, including LC retention time and MS2 spectra generated via DIA-NN (18, 20, 41).

Identified citrullinated peptides were classified as true positives (TP) if they matched the spiked synthetic peptides or as false positives (FP) if they did not. The minimal endogenous citrullination that may occur in the selected cell line is considered negligible due to the lack of detectable endogenous PADs (42) and citrullinated peptides in single-shot proteomics experiments.

Following this classification, we reported the TP and FP counts of PSMs in all samples from each search engine (Figure 2B; Supplementary Table S2). At a glance, MQ and MSFragger reported similar TP counts in the samples with proteome background, correlating well with the dilution ratios. However, MSFragger without MSBooster reported fewer FP counts compared to MQ and MSFragger with MSBooster. This result was evident in the precision metrics, where MSFragger exhibited much higher precision across the dilution range (Figure 2C; Supplementary Table S2). In biological samples without PAD enzyme overexpression and activation, e.g. human tissue proteomes, the abundance of citrullination falls between the sixth and seventh dilution or even lower (9) (Supplementary Table S3). In this range, the precision of the search engines drops below 50%, which may explain the high number of FPs observed in our previous study (9).

Another key observation is that while integrating MSBooster—a strategy that leverages peptide properties, including comparisons with predictive MS2 spectra—improves overall peptide identification, it is not universally effective for modified peptides. PTMs such as citrullination drastically alter fragmentation patterns, and comparisons with incorrect predictive spectra lead to higher FP counts in the evaluation dataset. Therefore, a model specifically trained on citrullinated peptides is necessary to exploit the potential of MS2 spectral features for accurate peptide identification.

### Training Prosit-Cit Model Using Synthetic Peptides from the ProteomeTools Library

Given the lack of an existing model to accurately predict the MS2 spectra of citrullinated peptides, we extended the Prosit deep learning framework to develop Prosit-Cit, a fine-tuned model specifically designed for this purpose. Prosit is a deep learning model recognized for its high accuracy in predicting the spectra of tryptic, non-tryptic, and isobaric-labeled peptides (17, 19, 21, 23, 24). Its accuracy largely stems from training on a vast spectral library, including ∼100 million high-quality MS2 spectra from 1.6 million synthetic peptide precursors within the ProteomeTools project (17, 19, 25).

To build Prosit-Cit, we leveraged high-quality LC-MS/MS datasets from ProteomeTools, including more than 30 million reference spectra from over 500,000 tryptic peptides, 242,000 non-tryptic peptides, and ∼2,500 synthetic citrullinated peptides with known citrullination sites (9, 25) (Figure 3A; Supplementary Table S4). It is worth noting that deamidated Asn and Gln residues are treated as Asp and Glu in the dataset, meaning that no specific deamidated peptides are required for training. Each peptide was measured and fragmented under various MS parameters, including CID and HCD fragmentation, and different HCD collision energies, forming the basis for the computational prediction of spectra and chromatographic retention times for both unmodified and citrullinated peptides. The detailed parameters can be found in ProteomeTools publication (25) and are also briefly described in the Methods section.

**Figure 3.**
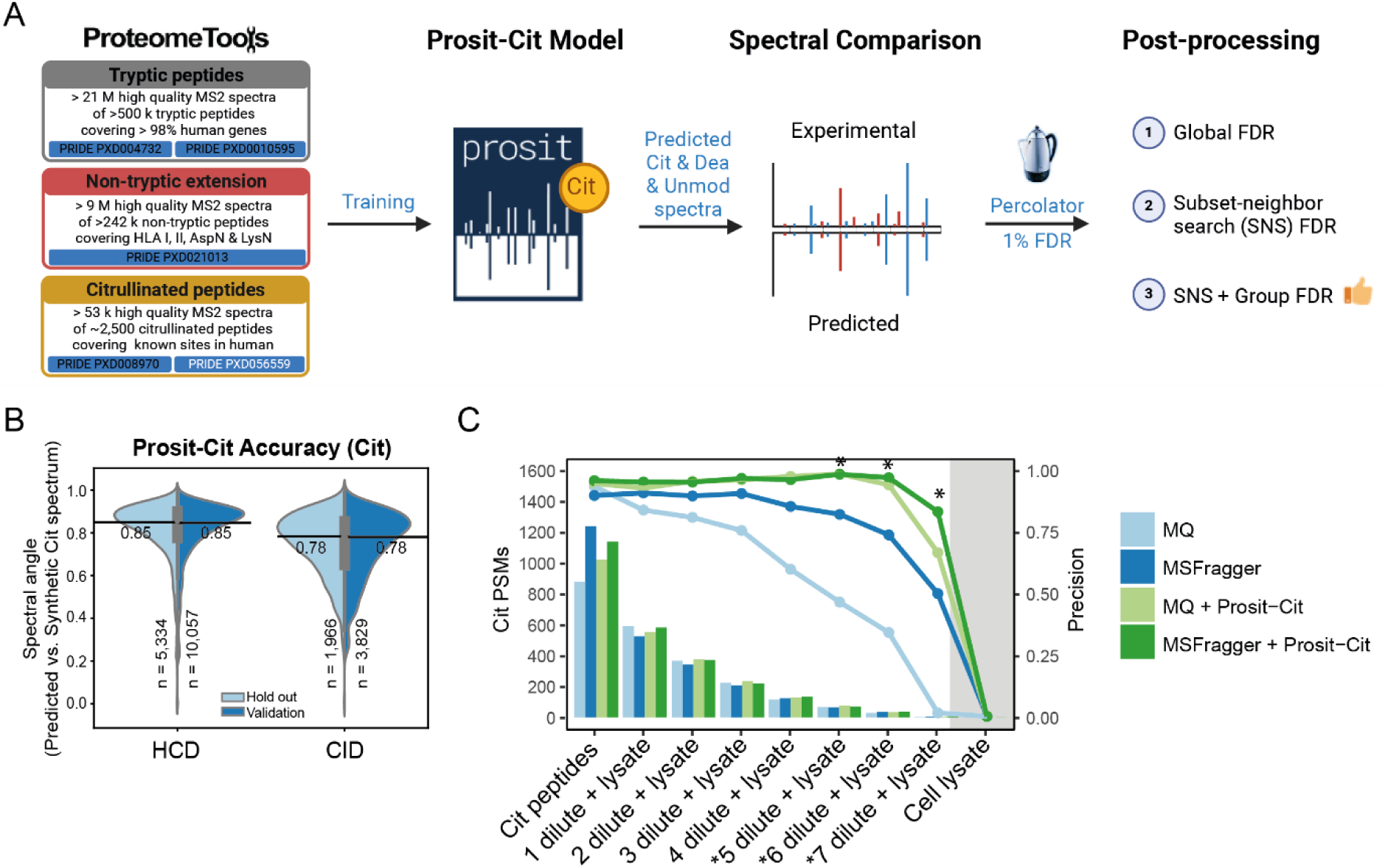
Prosit-Cit rescoring enhances the precision of citrullination identification. **A.** The deep learning framework Prosit-Cit was trained on data available prior to this study and data on citrullinated peptides with different collision energies and CID fragmentation type generated in this study (white). After rescoring, three post-processing approaches were examined. **B.** Beanplots comparing the prediction accuracy of the Prosit-Cit model between the holdout (light blue) and validation (dark blue) set in HCD and CID fragmentation types on citrullinated peptides. The black solid line and corresponding numbers indicate the median spectral angle for each distribution. The number of underlying spectra (n) is indicated at the bottom. **C.** Citrullinated PSMs and precision of identification using the SNS-Group FDR approach following database searching with MQ and MSFragger. Asterisk indicates the range of citrullination level in human tissue proteomes (9).

To evaluate the accuracy of Prosit-Cit for citrullinated peptides, we set aside ∼14,000 spectra from 205 citrullinated peptides as a validation set during training, and ∼7,300 spectra from 105 citrullinated peptides as a holdout set, with neither peptide sets contained in the training set. Despite the smaller dataset size, Prosit-Cit demonstrated strong performance, with median SA scores of 0.85 for HCD spectra and 0.78 for CID spectra (Figure 3B). The model also maintained high accuracy in predicting unmodified peptides, with median SA scores of 0.92 for HCD and 0.90 for CID spectra (Supplementary Figure S1A). The slightly lower median SA for citrullinated peptides was expected due to the limited size of the training set. Furthermore, the model showed high accuracy in predicting LC retention times for both citrullinated and unmodified peptides (ΔiRT 95% = 2.25-2.35 min for citrullinated peptides, ΔiRT 95% = 1.18-1.20 min for unmodified peptides, Pearson R ≥ 0.99, Supplementary Figure S1B).

### Prosit-Cit Rescoring Enhances Precision of Citrullination Identification

Given the high accuracy of the Prosit-Cit model in predicting citrullinated peptide spectra, we anticipated that integrating Prosit-Cit rescoring could greatly improve citrullination identification. To test this, we rescored our evaluation dataset results using Prosit-Cit via Oktoberfest (43), a Python package that facilitates spectral library generation and rescoring, with FDR control applied through Percolator (34).

However, FDR control for rare PTMs like citrullination is challenging, as the lower number of identifications can lead to skewed score distributions (14). To optimize FDR control for citrullination, we compared different post-processing methods with the conventional global FDR approach where all PSMs are used for FDR control (Supplementary Figure S1C&D; Supplementary Table S5). As the most common mistakes in database searching for citrullination identifications are misidentification of deamidation and the incorrect assignment of the monoisotopic peak (9, 12), a precursor mass with 0.98 Da increase could potentially be citrullinated, deamidated, or an unmodified peptide. To address this, we adapted a method similar to “subset-neighbor search” (SNS) (15). Here we took the subset of citrullination and deamidation identifications, and generated complementary deamidated/citrullinated/unmodified peptides as “neighbors” and selected the best-scoring version for FDR control. This method greatly improved precision, especially in MSFragger results in the lower dilution range (Supplementary Figure S1D; Supplementary Table S5). Considering the low physiological abundance of citrullination, we further refined precision by applying a second FDR control step similar to “group FDR” or “subset FDR” (44), focusing on citrullination and deamidation identifications only. This second step of FDR control decreased false positives and improved precision at the lowest dilution without sacrificing sensitivity (Supplementary Figure S1E; Supplementary Table S5).

We compared SNS-group FDR with Prosit rescoring to the results from MQ and MSFragger (Figure 3C). Prosit-Cit rescoring clearly enhanced precision in both MQ and MSFragger, with a more pronounced improvement in MQ results. Combining MSFragger and Prosit-Cit rescoring maintained a high precision (> 80%) across the wide range of dilutions. Based on these findings, we propose a data analysis workflow for citrullination identification that includes: (1) database searching with MSFragger due to shorter processing time, (2) generating neighbor peptides of citrullinated or deamidated peptides and rescoring identifications using the Prosit-Cit model via Koina (45), and (3) post-processing via Percolator with a two-step SNS-Group FDR control to maintain high precision and sensitivity. This pipeline (steps 2 and 3) is integrated into the Oktoberfest Python package as detailed in the Data availability section.

### Reanalyzing ten human tissue proteomes with MSFragger and Prosit-Cit boosts the identification of citrullination sites

Given that our validated pipeline gives high precision as well as sensitivity in the evaluation dataset, we hypothesized that this pipeline can greatly improve the precision and boost the identifications of existing proteomics datasets. To test this, we reanalyzed human tissue proteomes acquired by Wang et al. (27) that we had previously mined for protein citrullination (9). All the citrullination sites reported by the previous study were validated by manual inspection and/or synthetic reference spectral comparison. We selected ten tissues with the most citrullination sites (> 20 sites) and applied our validated pipeline. Despite using different search engines (Mascot vs. MSFragger) and post-processing steps (manual spectral inspection vs. automatic pipeline), our workflow successfully retrieved most of the citrullinated peptides and sites identified previously (Figure 4A; Supplementary Figure S2).

**Figure 4.**
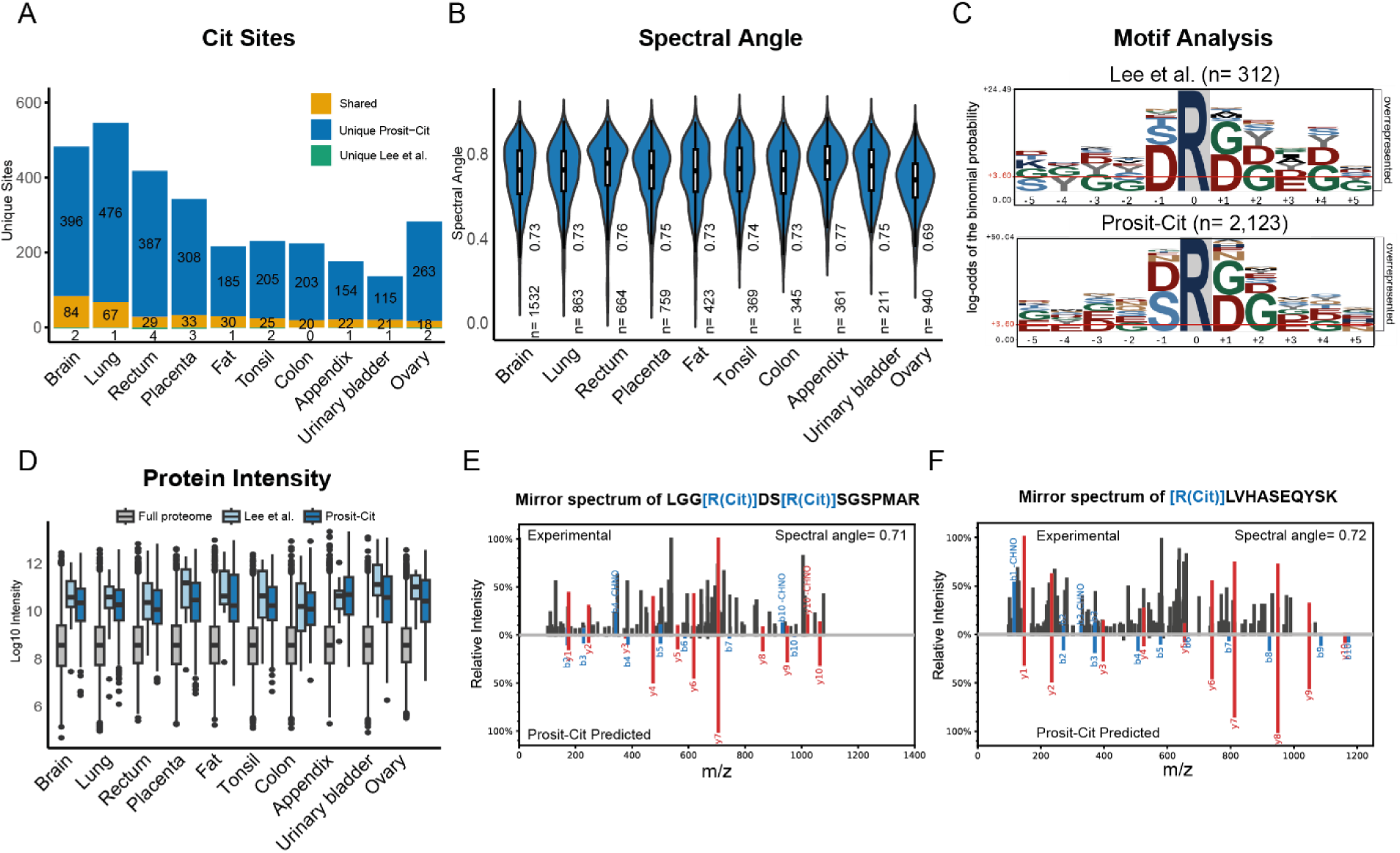
Reanalyzing ten human tissue proteomes with MSFragger and Prosit-Cit boosts the identification of citrullination sites. **A.** Identified citrullinated peptides in 10 human tissue proteomes acquired by Wang *et al.*(27). **B.** Spectral angle distributions in 10 human tissues. The numbers in the middle indicate the median spectral angle in each tissue. The number of underlying spectra (n=) is indicated at the bottom. **C.** Motif analysis based on the results from Lee *et al.* (9) and this study (Prosit-Cit). Citrullinated sites were shown as R at position zero. Numbers of sequences used are shown at the top (n). **D.** Protein intensity of the full proteome and the ones with citrullination identified in Lee *et al.* and Prosit-Cit. **E.** Mirror spectrum of the peptide from myelin basic protein in brain (top) compared to its predicted spectrum by Prosit-Cit (bottom). Spectral angle was shown on the top right. **F.** Mirror spectrum of the peptide from catenin delta-2 in brain (top) compared to its predicted spectrum by Prosit-Cit (bottom). Spectral angle is shown on the top right.

Furthermore, we observed a substantial increase in the identification of citrullinated peptides with our workflow (Figure 4A; Supplementary Table S6). This improvement is attributed not only to Prosit-Cit but also to MSFragger’s ability to identify 40-50% more PSMs compared to Mascot (Supplementary Table S7). When inspecting the spectral angles of PSMs of citrullinated peptides in each tissue, the median SA ranged from 0.69 to 0.76, indicating good similarity between predicted citrullinated spectra and experimental spectra (Figure 4B). While the SA values of the experimental data show a slight decrease compared to the validation and hold-out datasets, this is expected due to the optimal nature of synthetic peptide spectra compared to experimental data.

Additionally, the identified citrullinated peptides share a very similar motif to those found in our previous study (9) and a recently published proteome-wide citrullination study in cells (10) (Figure 4C). Given that we did not perform any enrichment for citrullinated proteins or peptides, it is expected that most proteins with identified citrullination sites were of high abundance. Examining the abundance of all proteins in the tissues revealed that our workflow is more sensitive, as it is able to identify citrullination sites in proteins of lower abundance compared to the previous study (Figure 4D). In cases where a peptide had more than one citrullination site (Figure 4E), spectral comparison provided confidence even with incomplete ion series. Similarly, for peptides with citrullination near the N-terminus (Figure 4F), where diagnostic ions were absent due to a lack of citrullination-containing y-ions, our workflow proved effective. In summary, our pipeline can increase sensitivity while maintaining the precision of citrullination identification in large-scale proteomics datasets.

### Exploring the citrullination landscape in Arabidopsis tissues

Having established a sensitive and precise pipeline for citrullinated peptide identification, we sought to explore citrullination in existing proteomics datasets. Although citrullination in humans is catalyzed by peptidylarginine deiminases (PADs) and PAD homologs are absent in non-vertebrate species (46), evidence suggests that enzymes with similar catalytic activity, e.g. hydrolase or amidinotransferase, can also citrullinate arginine residues (47). For instance, the bacterial enzyme peptidylarginine deiminase PPAD (48) and AGMATINE DEIMINASE/HYDROLASE (AIH; Q8GWW7; encoded by locus *At5g08170*) in Arabidopsis have been shown to catalyze citrullination (4). The later study identified about 20 citrullination sites in Arabidopsis root cell cultures.

To build on this, we applied our pipeline to a high-quality dataset of 30 Arabidopsis tissue proteomes (proteome depth > 12,000 proteins per tissue) (28). Our analysis identified 199 citrullination sites from 169 proteins across 30 tissues (Figure 5A; Supplementary Table S8), with the highest number of sites found in the hypocotyl, followed by the floral organs stamen and petals, while only one site was detected in the embryo or callus. Despite the lower number of citrullinated peptides identified compared to human proteomes, the median spectral angle (SA) scores were comparable, ranging from 0.61 to 0.75 across tissues (Figure 5B). Protein abundance across tissues showed that most proteins with identified citrullination sites were highly abundant, as expected (Supplementary Figure S3A). Grouping individual tissues into broader tissue groups revealed that flower tissues had the most citrullination sites, followed by the stem and leaf (Figure 5C; Supplementary Table S8). In addition, we identified nine citrullination sites present in at least six different tissues (20% of all tissues) (Figure 5D), including R46 in the UDP-GLUCOSE 6-DEHYDROGENASE 1 (UGD1; Q9FZE1; Supplementary Figure S3B) and R131 in the SMALL ribosomal subunit protein uS11c (RPS11; P56802; Supplementary Figure S3C). Although it is unclear whether these citrullination sites might alter the function of UGD1 or RPS11, both sites are located in functional binding domains (NAD- and RNA-binding, respectively) (49), suggesting possible regulatory roles. Additionally, seven proteins were found to have more than two citrullinated sites (Figure 5E), including the seed storage proteins CRUCIFERINA 1 (CRA1; P15455), CHLOROPLAST RNA BINDING (CRB; P15456), PLASTID-LIPID-ASSOCIATED PROTEIN 85 (PAP85; Q9LUJ7), and the elongation factor LOW EXPRESSION OF OSMOTICALLY RESPONSIVE GENES 1 (LOS1; Q9ASR1). LOS1 is a cold-induced translation elongation factor (type 2) that catalyzes the GTP-dependent ribosomal translocation step upon cold stress (50). Specifically, R350 of LOS1 is located at the GTP-binding domain’s C-terminus, suggesting potential GTP binding affinity alterations due to citrullination. The mirror spectra of LOS1 R350 from hypocotyl are shown in Figure 5F.

**Figure 5.**
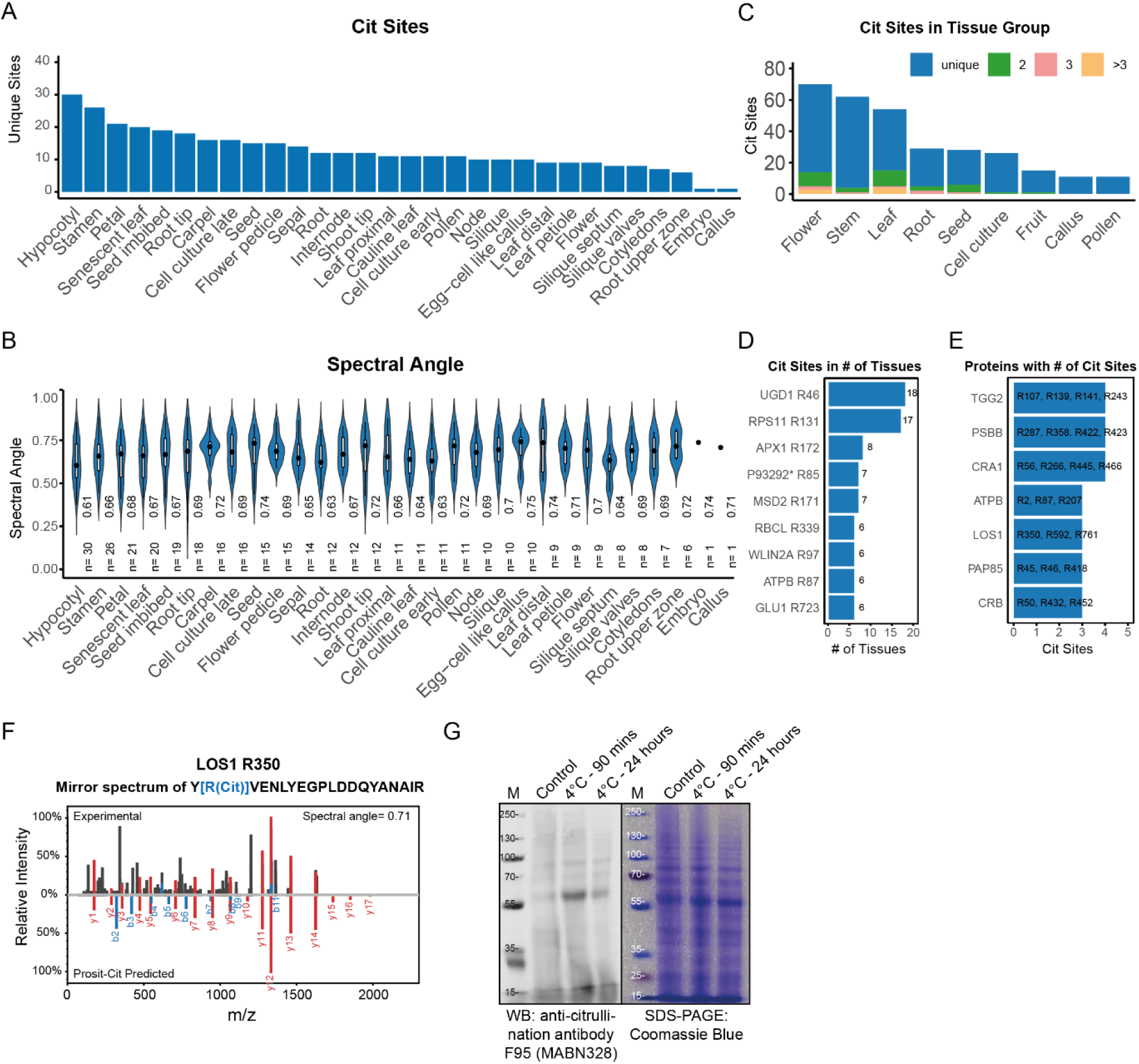
Exploring the citrullination landscape in Arabidopsis tissues. **A.** Identified citrullination sites in 30 Arabidopsis tissue proteomes acquired by Mergner *et al.* (28). **B.** Spectral angle distributions in 30 tissues. The numbers in the middle indicate the median spectral angle in each tissue. The number of underlying spectra (n=) is indicated at the bottom. **C.** Identified citrullination sites in tissue groups. The numbers of shared sites among tissues in the same tissue group is indicated by different colors. **D.** Citrullinated sites shared by more than six tissues. UGD1: UDP-GLUCOSE 6-DEHYDROGENASE 1; RPS11: SMALL RIBOSOMAL SUBUNIT PROTEIN US11C; APX1: L-ASCORBATE PEROXIDASE 1, CYTOSOLIC; P93292: Putative uncharacterized mitochondrial protein AtMg00280; MSD2: SUPEROXIDE DISMUTASE [MN] 2, MITOCHONDRIAL; RBCL: RIBULOSE BISPHOSPHATE CARBOXYLASE LARGE CHAIN; WLIN2A: LIM DOMAIN-CONTAINING PROTEIN WLIM2A; ATPB: ATP SYNTHASE SUBUNIT BETA, CHLOROPLASTIC; GLU1: FERREDOXIN-DEPENDENT GLUTAMATE SYNTHASE 1, CHLOROPLASTIC/MITOCHONDRIAL. **E.** Proteins with more than two citrullinated sites identified. TGG2: MYROSINASE 2; PSBB: PHOTOSYSTEM II CP47 REACTION CENTER PROTEIN; CRA1: 12S SEED STORAGE PROTEIN CRA1; ATPB: ATP SYNTHASE SUBUNIT BETA CHLOROPLASTIC; LOS1: ELONGATION FACTOR 2; PAP85: VICILIN-LIKE SEED STORAGE PROTEIN AT3G22640; CRB: 12S SEED STORAGE PROTEIN CRB. **F.** Mirror spectrum of citrullinated peptides of Elongation factor 2 (R350) from Hypocotyl (top) compared to its predicted spectrum by Prosit-Cit (bottom). **G.** Western blot of flower tissue using the anti-citrullination antibody F95 (MABN328) under short (90 mins) or long (24 hrs) cold stress at 4 °C. The control sample was incubated at 21°C.

Since a previous study had linked citrullination to cold stress (4), and this PTM was found to be most prevalent in floral tissues from our results, we investigated citrullination changes in flowers in response to cold. For this purpose, flowering Arabidopsis plants were exposed to cold stress (4°C) for 90 minutes or 24 hours, and the proteome of the flowers was resolved using LC-MS/MS. Despite achieving a good proteome coverage (12-13k proteins; Supplementary Table S9), we identified only six citrullination sites, with none of them consistently detected among all conditions (Supplementary Table S9). This may be due to the low citrullination level and the fact that no enrichment was performed before measurement. To verify this result, we assessed global citrullination changes using immunoblotting with the pan anti-citrulline antibody F95 (51) on these samples. The results showed an apparent increase in the citrullination signal after 90 minutes of cold stress, which returned to a baseline after 24 hours (Figure 5G). In summary, these observations suggest that citrullination may contribute to major cellular housekeeping processes, including protein translation, but also participate in environmental responses, such as those to cold stress, providing a basis for future studies that address the functional significance of this PTM in plants.

## Discussion

Citrullination is an important PTM with roles in both physiology and pathology, but its detection remains challenging due to the lack of efficient site-specific enrichment techniques and the complexity of identifying citrullinated peptides. Advances in mass spectrometry have opened new avenues for studying citrullination, yet scalable and precise data analysis pipelines for identifying citrullinated peptides in large proteomics datasets are still lacking. In this study, we developed a sensitive pipeline that addresses these challenges and enables high-confidence identification of citrullination sites.

We made two key technical improvements. First, we developed a dedicated Prosit model trained specifically to predict the MS2 spectra of citrullinated peptides. While a few existing models can predict spectra carrying modifications by shifting fragment ions, citrullination drastically alters fragmentation patterns, leading to decreased accuracy. Our results showed that MSBooster, designed to enhance peptide identification, increases both true and false positives, decreasing the precision (Figure 2B&C). Thus, training deep learning models using a tailored dataset was critical to achieving high prediction accuracy.

Second, we extended our rescoring pipeline to address the common challenge of distinguishing citrullinated from deamidated and unmodified peptides, as well as optimize the FDR control. Misidentification between citrullinated and deamidated peptides is particularly problematic due to their same mass shifts. To resolve this, we implemented a new approach similar to a “subset-neighbor search” (SNS) (15) strategy that generates alternative identifications (including deamidated and unmodified forms) for each potential citrullination site. This allowed us to evaluate competing hypotheses accurately. Additionally, we applied a second “group FDR” (44) approach that specifically focuses on citrullination and deamidation identifications, ensuring high precision in cases where the number of citrullination sites is low, as in Arabidopsis. This two-step approach effectively improves the reliability of identifications, even in datasets with low citrullination levels (Figure 3C).

Beyond technical improvements, our study also uncovered new biological insights. We identified approximately 200 citrullination sites across Arabidopsis tissues and validated changes in citrullination levels during cold stress through independent experiments. Despite the absence of a PAD homolog in plants, a previous study identified AIH as a potential citrullinating enzyme (4). However, our analysis found no clear correlation between AIH abundance and citrullination levels across tissues (Pearson correlation: −0.26; Supplementary Figure S3D). A possible explanation for the lack of a clear relationship between citrullination levels and AIH protein expression levels is that the enzymatic activity may be regulated under particular stress, and the abundance of it cannot indicate its activity. Interestingly, motif analysis revealed a citrullination pattern similar to humans (9, 10), with Gly and Asp preferred at the +1 position (Supplementary Figure S3E), suggesting a conserved motif across species. However, from a dataset with low citrullination site identifications, further research is needed to identify the enzymes responsible for citrullination in plants.

While we did not demonstrate it in this study, our Prosit-Cit model is able to predict spectra for non-tryptic citrullinated peptides and peptides fragmented by CID, which extends its applicability to a broader range of proteomic experiments. This is particularly relevant for mining datasets from immunopeptidomics, where citrullination was altered in tumor immunopeptidome (52).

Despite the sensitivity of our pipeline, most of the identified citrullination sites were found on highly abundant proteins (Figure 4D; Supplementary Figure S3A). This indicates that enrichment strategies will be critical for advancing citrullination studies, especially in systems with potentially lower modification levels, like Arabidopsis. Developing probes to selectively enrich citrullinated peptides before MS measurement will improve the depth of citrullinome analysis and enhance our understanding of its functional role.

We believe our pipeline opens new opportunities for the exploration of citrullination in biological research. By integrating this pipeline into the Oktoberfest Python package (43), we enable researchers to apply this method across existing proteomics datasets with high confidence. In the future, we anticipate that this workflow will drive deeper insights into the functional roles of citrullination in both plant and animal systems, contributing to a broader understanding of this PTM in various biological contexts.

## Abbreviations

Cit: citrullination/citrullinated
CID: collision-induced dissociation
Dea: Deamidation/deamidated
FDR: false discovery rate
FP: False Positive
HCD: higher-energy collision dissociation
IT: Ion trap
MS: Mass spectrometry
MS2: Tandem mass; MS/MS
OT: Orbitrap
PAD: peptidylarginine deiminases
PSM: peptide-spectrum match
SA: spectral angle
TP: True Positive

## Data availability

The mass spectrometry proteomics data have been deposited to the ProteomeXchange Consortium (http://proteomecentral.proteomexchange.org) via the PRIDE partner repository (53). Part of the reference spectra (HCD) of synthetic citrullinated peptides have been published previously (9) and are available under the dataset identifier PXD008970. The raw data, search and rescoring results of the evaluation and Arabidopsis flower dataset are deposited under the dataset identifier PXD056560. The raw data and search results of the ProteomeTools synthetic citrullinated peptides from CID fragmentation and different collision energies are deposited under the dataset identifier PXD056559. Annotated spectra accompanied by metadata are archived as parquet files in Zenodo (https://zenodo.org/records/13856705). The tutorial Jupyter notebook is available on GitHub (https://github.com/wilhelm-lab/oktoberfest/tree/development/tutorials).

## Author contributions

C.-Y.L. and M.W. conceived and jointly supervised the study. C.-Y.L., M.W., W.G., G.D., B.P. and R.M.G. designed experiments. W.G., R.M.G., S.L., E.R., and G.D. performed experiments. C.-Y.L, W.G., R.M.G., and M.W. analyzed data. C.-Y.L., W.G., and R.M.G. wrote the manuscript with final input from all authors.

## Acknowledgments

The authors would like to express their gratitude to Prof. Dr. Bernhard Küster, Dr. Christina Ludwig, and all members of the Lee Lab, the Bavarian Center for Biomolecular Mass Spectrometry (BayBioMS), and the Chair of Proteomics and Bioanalytics for their valuable assistance and insightful discussions. Special thanks to Prof. Dr. Bernhard Küster for generously providing access to the ProteomeTools resource. This work was partly funded by the Federal Ministry of Education and Research (FKZ161L0215; YIG-SysNS), European Union’s Horizon 2020 Program under Grant Agreement 823839 (H2020-INFRAIA-2018-1; EPIC-XS) and ERC Starting Grant (grant number 101077037). The Orbitrap Fusion Lumos and HF-X mass spectrometer used in this study were funded in part by the German Research Foundation (DFG-INST 95/1436-1 FUGG & DFG-INST 95/1435-1 FUGG). The graphical abstract and illustrations were created with BioRender.com.

## Supplementary Figures

**Figure S1.**
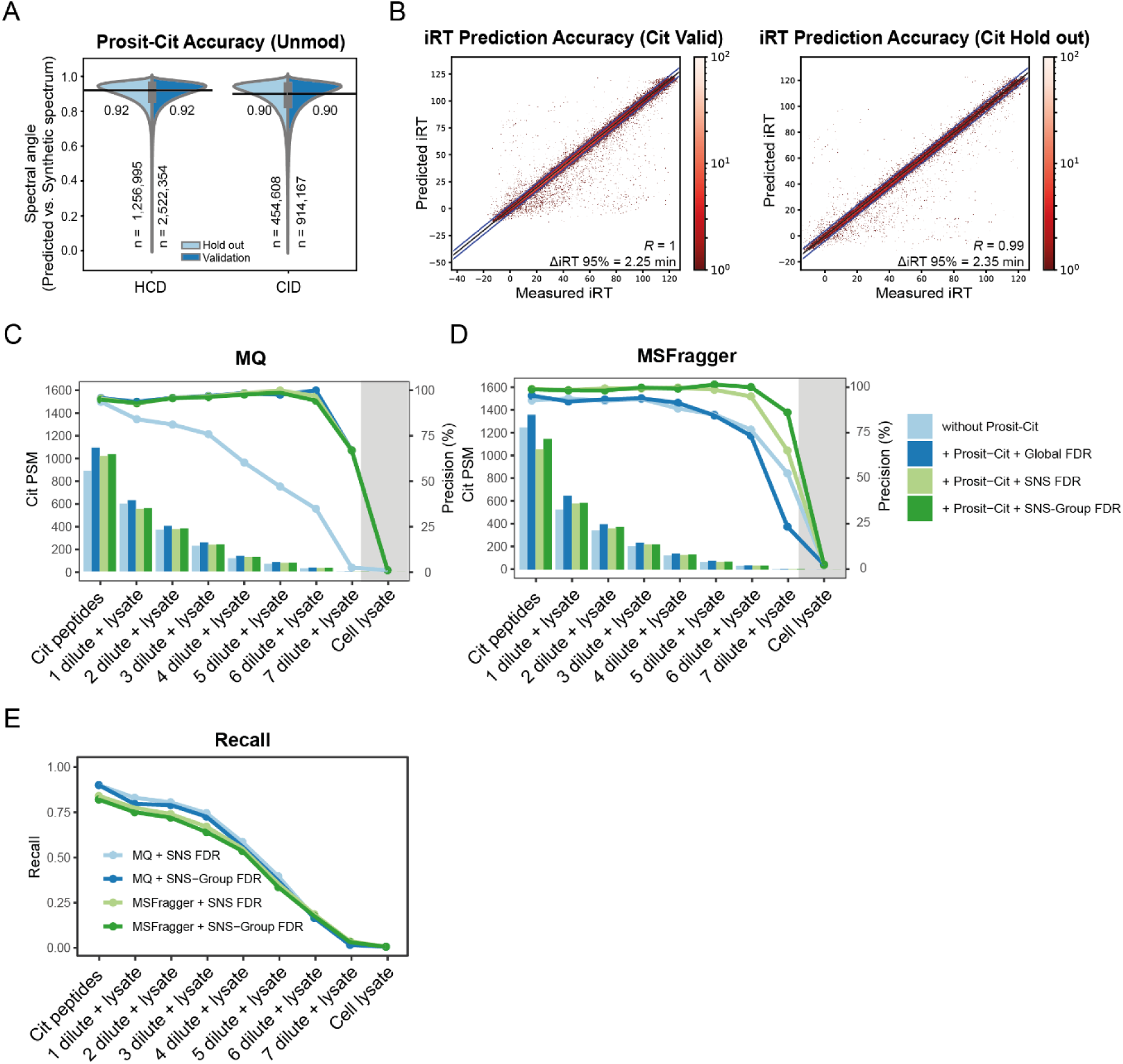
Prosit-Cit accuracy and influences of four post-processing methods. **A.** Beanplots of the prediction accuracy of Prosit-Cit model between the holdout (light blue) and validation (dark blue) set in HCD and CID fragmentation types on unmodified peptides. The black solid line and corresponding numbers indicate the median spectral angle for each distribution. The number of underlying spectra (n) is indicated at the bottom. **B.** Scatter plot of predicted indexed retention times (iRT) with the Prosit-Cit model compared to experimentally determined iRTs of citrullinated peptides in the holdout and validation set. The delta iRT necessary to encompass 95% of the peptide (ΔiRT95%) and the Pearson’s R is indicated in the bottom. **C-D.** Citrullinated PSMs and precision of identification using different post-processing approaches following database searching with MaxQuant (C) and MSFragger (D). **E.** Recall of identified citrullinated peptides using different post-processing approaches. The TP count was divided by the total numbers of synthetic citrullinated peptides spiked-in.

**Figure S2.**
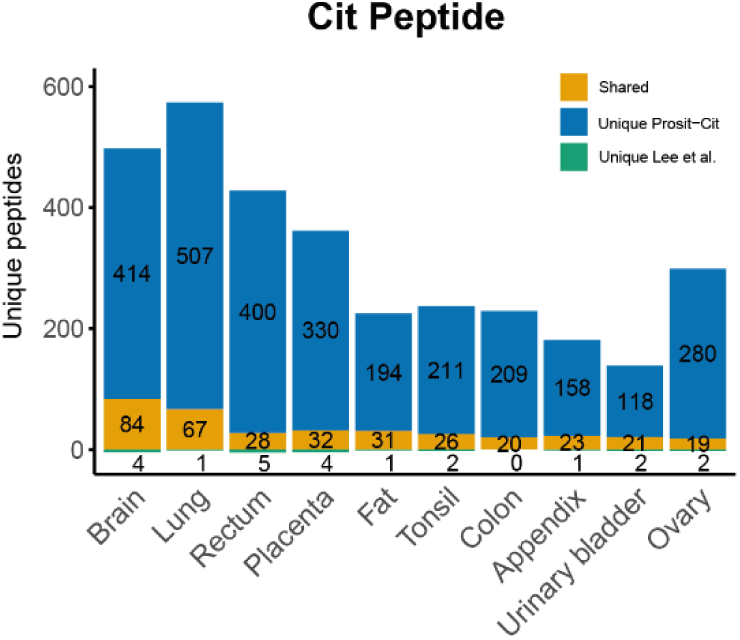
Identified citrullinated peptides in 10 human tissue proteomes in Lee *et al.* (9) and this study (Prosit-Cit).

**Figure S3.**
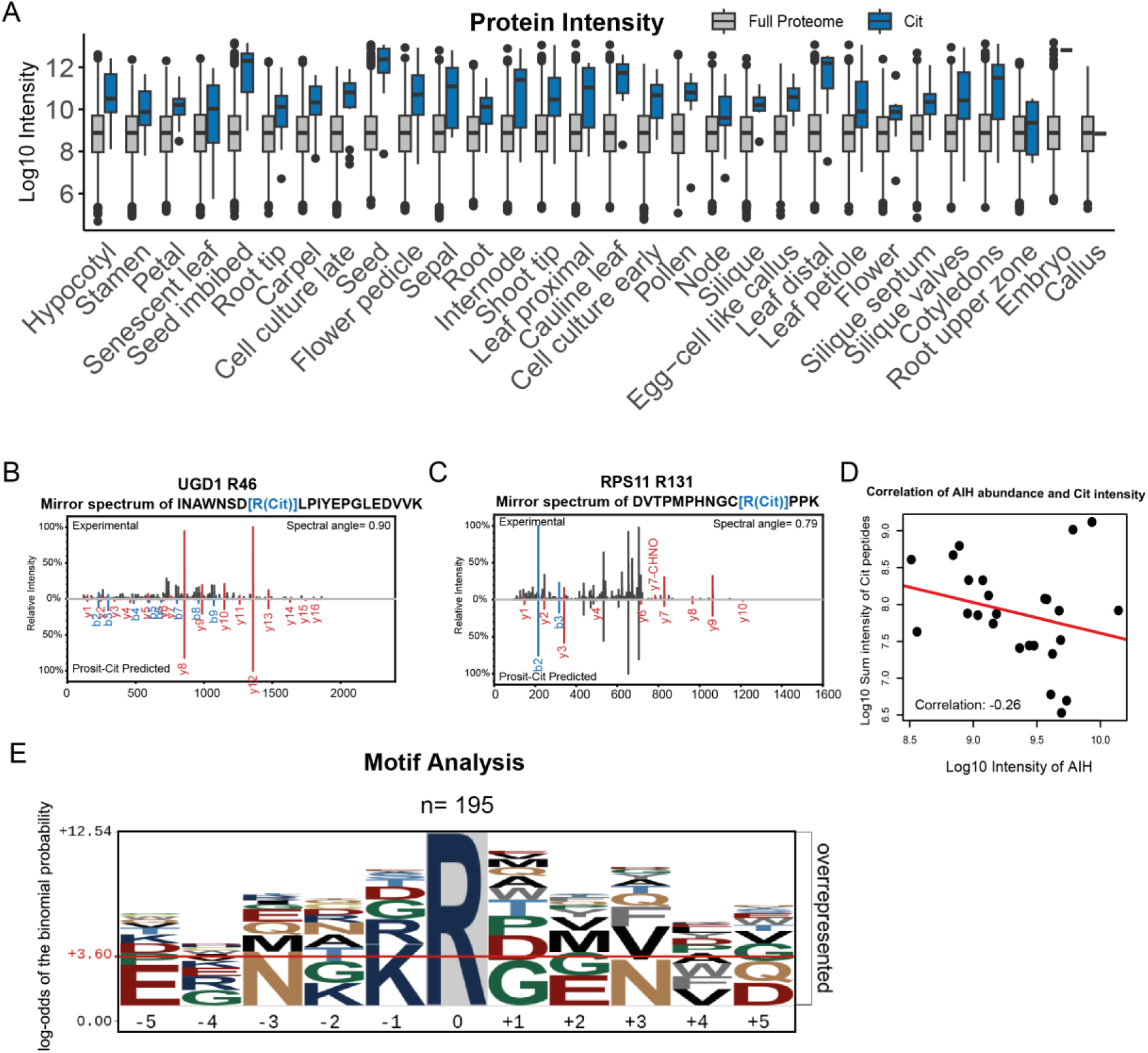
Exploring the citrullination landscape in Arabidopsis tissues. **A.** Protein intensity of the full proteome and proteins with citrullination sites identified. **B.** Mirror spectrum of citrullinated peptides of UGD1 R46 (Q9FZE1; UDP-GLUCOSE 6-DEHYDROGENASE 1) from flower pedicle (top) compared to its predicted spectrum by Prosit-Cit (bottom). **C.** Mirror spectrum of citrullinated peptides of RPS11 R131 (P56802; SMALL RIBOSOMAL SUBUNIT PROTEIN US11C from cotyledons (top) compared to its predicted spectrum by Prosit-Cit (bottom). **D.** Correlation of the abundance of AGMATINE DEIMINASE/HYDROLASE (AIH) with the sum intensity of citrullinated peptides across tissues. Pearson’s correlation is shown. **E.** Motif analysis of citrullination sites. Citrullination sites are shown as R at position zero. Numbers of sequences used are shown at the top.

## References

1. Lundberg, K., Nijenhuis, S., Vossenaar, E. R., Palmblad, K., van Venrooij, W. J., Klareskog, L., Zendman, A. J., and Harris, H. E. (2005) Citrullinated proteins have increased immunogenicity and arthritogenicity and their presence in arthritic joints correlates with disease severity. Arthritis Res Ther 7, R458–467

2. Christophorou, M. A. (2022) The virtues and vices of protein citrullination. R Soc Open Sci 9, 220125

3. Gyorgy, B., Toth, E., Tarcsa, E., Falus, A., and Buzas, E. I. (2006) Citrullination: a posttranslational modification in health and disease. Int J Biochem Cell Biol 38, 1662–1677

4. Marondedze, C., Elia, G., Thomas, L., Wong, A., and Gehring, C. (2021) Citrullination of Proteins as a Specific Response Mechanism in Plants. Front Plant Sci 12, 638392

5. Doll, S., and Burlingame, A. L. (2015) Mass spectrometry-based detection and assignment of protein posttranslational modifications. ACS Chem Biol 10, 63–71

6. Zhao, Y., and Jensen, O. N. (2009) Modification-specific proteomics: strategies for characterization of post-translational modifications using enrichment techniques. Proteomics 9, 4632–4641

7. Tutturen, A. E., Fleckenstein, B., and de Souza, G. A. (2014) Assessing the citrullinome in rheumatoid arthritis synovial fluid with and without enrichment of citrullinated peptides. J Proteome Res 13, 2867–2873

8. Lewallen, D. M., Bicker, K. L., Subramanian, V., Clancy, K. W., Slade, D. J., Martell, J., Dreyton, C. J., Sokolove, J., Weerapana, E., and Thompson, P. R. (2015) Chemical Proteomic Platform To Identify Citrullinated Proteins. ACS Chem Biol 10, 2520–2528

9. Lee, C. Y., Wang, D., Wilhelm, M., Zolg, D. P., Schmidt, T., Schnatbaum, K., Reimer, U., Ponten, F., Uhlen, M., Hahne, H., and Kuster, B. (2018) Mining the Human Tissue Proteome for Protein Citrullination. Mol Cell Proteomics 17, 1378–1391

10. Rebak, A. S., Hendriks, I. A., Elsborg, J. D., Buch-Larsen, S. C., Nielsen, C. H., Terslev, L., Kirsch, R., Damgaard, D., Doncheva, N. T., Lennartsson, C., Rykaer, M., Jensen, L. J., Christophorou, M. A., and Nielsen, M. L. (2024) A quantitative and site-specific atlas of the citrullinome reveals widespread existence of citrullination and insights into PADI4 substrates. Nat Struct Mol Biol 31, 977–995

11. Di Palma, S., Hennrich, M. L., Heck, A. J., and Mohammed, S. (2012) Recent advances in peptide separation by multidimensional liquid chromatography for proteome analysis. J Proteomics 75, 3791–3813

12. Wang, X., Swensen, A. C., Zhang, T., Piehowski, P. D., Gaffrey, M. J., Monroe, M. E., Zhu, Y., Dong, H., and Qian, W. J. (2020) Accurate Identification of Deamidation and Citrullination from Global Shotgun Proteomics Data Using a Dual-Search Delta Score Strategy. J Proteome Res 19, 1863–1872

13. Maurais, A. J., Salinger, A. J., Tobin, M., Shaffer, S. A., Weerapana, E., and Thompson, P. R. (2021) A Streamlined Data Analysis Pipeline for the Identification of Sites of Citrullination. Biochemistry 60, 2902–2914

14. Fu, Y., and Qian, X. (2014) Transferred subgroup false discovery rate for rare post-translational modifications detected by mass spectrometry. Mol Cell Proteomics 13, 1359–1368

15. Lin, A., Plubell, D. L., Keich, U., and Noble, W. S. (2021) Accurately Assigning Peptides to Spectra When Only a Subset of Peptides Are Relevant. J Proteome Res 20, 4153–4164

16. Shortreed, M. R., Wenger, C. D., Frey, B. L., Sheynkman, G. M., Scalf, M., Keller, M. P., Attie, A. D., and Smith, L. M. (2015) Global Identification of Protein Post-translational Modifications in a Single-Pass Database Search. J Proteome Res 14, 4714–4720

17. Gessulat, S., Schmidt, T., Zolg, D. P., Samaras, P., Schnatbaum, K., Zerweck, J., Knaute, T., Rechenberger, J., Delanghe, B., Huhmer, A., Reimer, U., Ehrlich, H. C., Aiche, S., Kuster, B., and Wilhelm, M. (2019) Prosit: proteome-wide prediction of peptide tandem mass spectra by deep learning. Nat Methods 16, 509–518

18. Demichev, V., Messner, C. B., Vernardis, S. I., Lilley, K. S., and Ralser, M. (2020) DIA-NN: neural networks and interference correction enable deep proteome coverage in high throughput. Nat Methods 17, 41–44

19. Wilhelm, M., Zolg, D. P., Graber, M., Gessulat, S., Schmidt, T., Schnatbaum, K., Schwencke-Westphal, C., Seifert, P., de Andrade Kratzig, N., Zerweck, J., Knaute, T., Braunlein, E., Samaras, P., Lautenbacher, L., Klaeger, S., Wenschuh, H., Rad, R., Delanghe, B., Huhmer, A., Carr, S. A., Clauser, K. R., Krackhardt, A. M., Reimer, U., and Kuster, B. (2021) Deep learning boosts sensitivity of mass spectrometry-based immunopeptidomics. Nat Commun 12, 3346

20. Yang, K. L., Yu, F., Teo, G. C., Li, K., Demichev, V., Ralser, M., and Nesvizhskii, A. I. (2023) MSBooster: improving peptide identification rates using deep learning-based features. Nat Commun 14, 4539

21. Adams, C., Gabriel, W., Laukens, K., Picciani, M., Wilhelm, M., Bittremieux, W., and Boonen, K. (2024) Fragment ion intensity prediction improves the identification rate of non-tryptic peptides in timsTOF. Nat Commun 15, 3956

22. Yi, X., Wen, B., Ji, S., Saltzman, A. B., Jaehnig, E. J., Lei, J. T., Gao, Q., and Zhang, B. (2024) Deep Learning Prediction Boosts Phosphoproteomics-Based Discoveries Through Improved Phosphopeptide Identification. Mol Cell Proteomics 23, 100707

23. Gabriel, W., Giurcoiu, V., Lautenbacher, L., and Wilhelm, M. (2022) Predicting fragment intensities and retention time of iTRAQ- and TMTPro-labeled peptides with Prosit-TMT. Proteomics 22, e2100257

24. Gabriel, W., The, M., Zolg, D. P., Bayer, F. P., Shouman, O., Lautenbacher, L., Schnatbaum, K., Zerweck, J., Knaute, T., Delanghe, B., Huhmer, A., Wenschuh, H., Reimer, U., Medard, G., Kuster, B., and Wilhelm, M. (2022) Prosit-TMT: Deep Learning Boosts Identification of TMT-Labeled Peptides. Anal Chem 94, 7181–7190

25. Zolg, D. P., Wilhelm, M., Schmidt, T., Medard, G., Zerweck, J., Knaute, T., Wenschuh, H., Reimer, U., Schnatbaum, K., and Kuster, B. (2018) ProteomeTools: Systematic Characterization of 21 Post-translational Protein Modifications by Liquid Chromatography Tandem Mass Spectrometry (LC-MS/MS) Using Synthetic Peptides. Mol Cell Proteomics 17, 1850–1863

26. Zolg, D. P., Wilhelm, M., Schnatbaum, K., Zerweck, J., Knaute, T., Delanghe, B., Bailey, D. J., Gessulat, S., Ehrlich, H. C., Weininger, M., Yu, P., Schlegl, J., Kramer, K., Schmidt, T., Kusebauch, U., Deutsch, E. W., Aebersold, R., Moritz, R. L., Wenschuh, H., Moehring, T., Aiche, S., Huhmer, A., Reimer, U., and Kuster, B. (2017) Building ProteomeTools based on a complete synthetic human proteome. Nat Methods 14, 259–262

27. Wang, D., Eraslan, B., Wieland, T., Hallstrom, B., Hopf, T., Zolg, D. P., Zecha, J., Asplund, A., Li, L. H., Meng, C., Frejno, M., Schmidt, T., Schnatbaum, K., Wilhelm, M., Ponten, F., Uhlen, M., Gagneur, J., Hahne, H., and Kuster, B. (2019) A deep proteome and transcriptome abundance atlas of 29 healthy human tissues. Mol Syst Biol 15, e8503

28. Mergner, J., Frejno, M., List, M., Papacek, M., Chen, X., Chaudhary, A., Samaras, P., Richter, S., Shikata, H., Messerer, M., Lang, D., Altmann, S., Cyprys, P., Zolg, D. P., Mathieson, T., Bantscheff, M., Hazarika, R. R., Schmidt, T., Dawid, C., Dunkel, A., Hofmann, T., Sprunck, S., Falter-Braun, P., Johannes, F., Mayer, K. F. X., Jurgens, G., Wilhelm, M., Baumbach, J., Grill, E., Schneitz, K., Schwechheimer, C., and Kuster, B. (2020) Mass-spectrometry-based draft of the Arabidopsis proteome. Nature 579, 409–414

29. Wenschuh, H., Volkmer-Engert, R., Schmidt, M., Schulz, M., Schneider-Mergener, J., and Reineke, U. (2000) Coherent membrane supports for parallel microsynthesis and screening of bioactive peptides. Biopolymers 55, 188–206

30. Hughes, C. S., Moggridge, S., Muller, T., Sorensen, P. H., Morin, G. B., and Krijgsveld, J. (2019) Single-pot, solid-phase-enhanced sample preparation for proteomics experiments. Nat Protoc 14, 68–85

31. Cox, J., and Mann, M. (2008) MaxQuant enables high peptide identification rates, individualized p.p.b.-range mass accuracies and proteome-wide protein quantification. Nat Biotechnol 26, 1367–1372

32. Kong, A. T., Leprevost, F. V., Avtonomov, D. M., Mellacheruvu, D., and Nesvizhskii, A. I. (2017) MSFragger: ultrafast and comprehensive peptide identification in mass spectrometry-based proteomics. Nat Methods 14, 513–520

33. Shouman, O., Gabriel, W., Giurcoiu, V.-G., Sternlicht, V., and Wilhelm, M. (2022) PROSPECT: Labeled Tandem Mass Spectrometry Dataset for Machine Learning in Proteomics. In: Koyejo, S., Mohamed, S., Agarwal, A., Belgrave, D., Cho, K., and Oh, A., eds., pp. 32882--32896

34. Kall, L., Canterbury, J. D., Weston, J., Noble, W. S., and MacCoss, M. J. (2007) Semi-supervised learning for peptide identification from shotgun proteomics datasets. Nat Methods 4, 923–925

35. O’Shea, J. P., Chou, M. F., Quader, S. A., Ryan, J. K., Church, G. M., and Schwartz, D. (2013) pLogo: a probabilistic approach to visualizing sequence motifs. Nat Methods 10, 1211–1212

36. Brajkovic, S., Rugen, N., Agius, C., Berner, N., Eckert, S., Sakhteman, A., Schwechheimer, C., and Kuster, B. (2023) Getting Ready for Large-Scale Proteomics in Crop Plants. Nutrients 15

37. Ruprecht, B., Zecha, J., Zolg, D. P., and Kuster, B. (2017) High pH Reversed-Phase Micro-Columns for Simple, Sensitive, and Efficient Fractionation of Proteome and (TMT labeled) Phosphoproteome Digests. Methods Mol Biol 1550, 83–98

38. Jin, Z., Fu, Z., Yang, J., Troncosco, J., Everett, A. D., and Van Eyk, J. E. (2013) Identification and characterization of citrulline-modified brain proteins by combining HCD and CID fragmentation. Proteomics 13, 2682–2691

39. Steckel, A., and Schlosser, G. (2019) Citrulline Effect Is a Characteristic Feature of Deiminated Peptides in Tandem Mass Spectrometry. J Am Soc Mass Spectrom 30, 1586–1591

40. Wan, K. X., Vidavsky, I., and Gross, M. L. (2002) Comparing similar spectra: from similarity index to spectral contrast angle. J Am Soc Mass Spectrom 13, 85–88

41. Demichev, V., Szyrwiel, L., Yu, F., Teo, G. C., Rosenberger, G., Niewienda, A., Ludwig, D., Decker, J., Kaspar-Schoenefeld, S., Lilley, K. S., Mulleder, M., Nesvizhskii, A. I., and Ralser, M. (2022) dia-PASEF data analysis using FragPipe and DIA-NN for deep proteomics of low sample amounts. Nat Commun 13, 3944

42. Lee, C. Y., The, M., Meng, C., Bayer, F. P., Putzker, K., Muller, J., Streubel, J., Woortman, J., Sakhteman, A., Resch, M., Schneider, A., Wilhelm, S., and Kuster, B. (2024) Illuminating phenotypic drug responses of sarcoma cells to kinase inhibitors by phosphoproteomics. Mol Syst Biol 20, 28–55

43. Picciani, M., Gabriel, W., Giurcoiu, V. G., Shouman, O., Hamood, F., Lautenbacher, L., Jensen, C. B., Muller, J., Kalhor, M., Soleymaniniya, A., Kuster, B., The, M., and Wilhelm, M. (2024) Oktoberfest: Open-source spectral library generation and rescoring pipeline based on Prosit. Proteomics 24, e2300112

44. Yi, X., Gong, F., and Fu, Y. (2020) Transfer posterior error probability estimation for peptide identification. BMC Bioinformatics 21, 173

45. Lautenbacher, L., Yang, K. L., Kockmann, T., Panse, C., Chambers, M., Kahl, E., Yu, F., Gabriel, W., Bold, D., Schmidt, T., Li, K., MacLean, B., Nesvizhskii, A. I., and Wilhelm, M. (2024) Koina: Democratizing machine learning for proteomics research. bioRxiv

46. Cummings, T. F. M., Gori, K., Sanchez-Pulido, L., Gavriilidis, G., Moi, D., Wilson, A. R., Murchison, E., Dessimoz, C., Ponting, C. P., and Christophorou, M. A. (2022) Citrullination Was Introduced into Animals by Horizontal Gene Transfer from Cyanobacteria. Mol Biol Evol 39

47. Linsky, T., and Fast, W. (2010) Mechanistic similarity and diversity among the guanidine-modifying members of the pentein superfamily. Biochim Biophys Acta 1804, 1943–1953

48. Maresz, K. J., Hellvard, A., Sroka, A., Adamowicz, K., Bielecka, E., Koziel, J., Gawron, K., Mizgalska, D., Marcinska, K. A., Benedyk, M., Pyrc, K., Quirke, A. M., Jonsson, R., Alzabin, S., Venables, P. J., Nguyen, K. A., Mydel, P., and Potempa, J. (2013) Porphyromonas gingivalis facilitates the development and progression of destructive arthritis through its unique bacterial peptidylarginine deiminase (PAD). PLoS Pathog 9, e1003627

49. Mistry, J., Chuguransky, S., Williams, L., Qureshi, M., Salazar, G. A., Sonnhammer, E. L. L., Tosatto, S. C. E., Paladin, L., Raj, S., Richardson, L. J., Finn, R. D., and Bateman, A. (2021) Pfam: The protein families database in 2021. Nucleic Acids Res 49, D412–D419

50. Guo, Y., Xiong, L., Ishitani, M., and Zhu, J. K. (2002) An Arabidopsis mutation in translation elongation factor 2 causes superinduction of CBF/DREB1 transcription factor genes but blocks the induction of their downstream targets under low temperatures. Proc Natl Acad Sci U S A 99, 7786–7791

51. Nicholas, A. P., and Whitaker, J. N. (2002) Preparation of a monoclonal antibody to citrullinated epitopes: its characterization and some applications to immunohistochemistry in human brain. Glia 37, 328–336

52. Kacen, A., Javitt, A., Kramer, M. P., Morgenstern, D., Tsaban, T., Shmueli, M. D., Teo, G. C., da Veiga Leprevost, F., Barnea, E., Yu, F., Admon, A., Eisenbach, L., Samuels, Y., Schueler-Furman, O., Levin, Y., Nesvizhskii, A. I., and Merbl, Y. (2023) Post-translational modifications reshape the antigenic landscape of the MHC I immunopeptidome in tumors. Nat Biotechnol 41, 239–251

53. Perez-Riverol, Y., Bai, J., Bandla, C., Garcia-Seisdedos, D., Hewapathirana, S., Kamatchinathan, S., Kundu, D. J., Prakash, A., Frericks-Zipper, A., Eisenacher, M., Walzer, M., Wang, S., Brazma, A., and Vizcaino, J. A. (2022) The PRIDE database resources in 2022: a hub for mass spectrometry-based proteomics evidences. Nucleic Acids Res 50, D543–D552

